# Simultaneous selection for yield and stability in cassava (*Manihot esculenta Crantz*) under optimum condition in Nigeria

**DOI:** 10.64898/2026.01.17.700121

**Authors:** G. O. Oyebode, M.O. Oayiwola

## Abstract

High yield and stability of performance across environments are required for success in agricultural systems. Selection of high yielding and stable genotypes is always hindered by Genotype by environment interaction (GEI). Hence plant breeders have proposed diverse methods to help increase their efficiency of selection in multi environment yield trials. The aim of this experiment was to identify and select high yielding and stable cassava genotypes using to GGE biplot and Yield selection index (YSi) techniques and to determine the efficiency of the techniques. 24 hybrids and 2 checks were tested across five locations in Nigeria using the randomized complete block design with 4 replications. ANOVA provided clear evidence of GEI (P<0.001) for dry root yield. 23 of the hybrids showed 10% to 83% economic improvement over the best check variety. GGE biplot identified genotypes TMS-IBA090574, TMS-IBA090521, TMS-IBA090590 and TMS-IBA090576 as high yielding and stable, while YSi selected 14 genotypes TMS-IBA090454, TMS-IBA090504, TMS-IBA090506, TMS-IBA090509, TMS-IBA090521, TMS-IBA090523, TMS-IBA090536, TMS-IBA090564, TMS-IBA090574, TMS-IBA090576, TMS-IBA090581, TMS-IBA090590, TMS-IBA090597 and TMS-IBA090609. Six of these genotypes (TMS-IBA090523, TMS-IBA090536, TMS-IBA090574, TMS-IBA090581, TMS-IBA090506, TMS-IBA090521) were marked as unstable by the Shukla’s stability variance which indicates a weakness of YSi. The two techniques identified TMS-IBA090590 and TMS-IBA090576 as high yielding and stable genotypes. These two clones performed above average in at least four of the five locations used for evaluation. The genotypes jointly selected by both methods may therefore be released to increase the productivity of the crop in Nigeria and comparable agro ecology.

## Introduction

Growing world population demands continuous food production to meet human food and nutritional needs. The need to combat hunger is clearly reflected in the Sustainable Development Goals (SDGs) of the UN, whose target is to eradicate hunger by 2030 (FAO, 2015). Achieving food and nutritional security is closely linked to increased production and availability of crops that are key food sources to the populace.

In sub-Saharan Africa (SSA) roots and tubers contribute a substantial amount of the required calories; they are highly nutritional, cheap, and therefore form essential constituent of human diets in the region. Among roots and tubers that serve as food, cassava (*Manihot esculenta* Crantz) is the most important. It is well adapted to most agro-ecologies in SSA, thrives and produces reasonable yields where most crops fail, thereby fitting as a food security crop and important source of dietary energy (FAO, 2011). Cassava forms an essential part of the diet for more than half a billion people and provides livelihood for millions of farmers, processors and traders worldwide (FAO, 2015). FAO 2015 food outlook report put global production of cassava at 202.6 million tonnes on 18.5 million hectares, with Nigeria, the largest producer, accounting for 57 million tonnes from a cultivated area of 3.8 million hectares.

The demand for cassava is on the rise particularly with more industries utilizing the roots as raw materials. Increasing cassava yields therefore becomes pertinent so that industrial usage does not affect its availability as a food source. However, cassava is mostly grown by smallholder farmers who rely greatly on landraces that are susceptible to pest and diseases and consequently low yielding. The development of high yielding and superior genotypes therefore becomes imperative. Identification and release of such genotypes is however notoriously hindered by Genotype x environment interaction (GEI).

GEI refers to the variation in the response of genotypes to varied environmental (season, location, years, crop management practices etc) conditions such that genotypes with desirable performance under a growing condition may be poor in another environment (Ceccarelli, 1996). However, the understanding of GEI patterns from GEI analyses has contributed tremendously to cultivar development. For instance, GEI has necessitated multi-environment trials (METS) from which breeders have identified genotypes that are adapted to a particular environment and those superior across several growing regions (Annicchiarico et al., 1996). Test environments have also been categorised and representative sites identified, thereby eliminating unnecessary test locations and reducing the cost of cultivar evaluation (Yan et al. 2001). The direct implication of a significant GEI is the need to select genotypes based on consistent positive performance over a wide range of environments. The selection of high yielding but unstable genotypes in a breeding program or commercial farm will lead to devastating results.

Plant breeders have thus developed several statistical techniques, including Genotype+Genotype x environment (GGE) biplot by Yan (2001) and Yield Stability Index (YSi), to aid decision on the superiority or desirability of genotypes and exploit the potentials of GEI. The efficiency of GGE biplot have been compared with other techniques by various workers for cassava (Aina et al., 2009; Agyeman et al., 2015) for more reliable decision. GGE biplot is the most recent and sophisticated among the proposed techniques (Yan et al., 2007) while YSi is superior among methods that integrate yield with stability rating (Waldron et al., 2002; Olayiwola and Ariyo, 2013). The aim of the study were to identify high yielding and stable cassava clones to be released for cultivation and compare two procedures for relative efficiency in the selection of stable and high yielding cassava clones for dry root yield. Information from this work will be crucial to efforts aimed at increasing cassava yield on farmers’ field to meet local and international demands.

## Materials and methods

The genotypes used for the experiment comprised of 24 F_1_ hybrids with high resistance to Cassava mosaic disease (CMD) and high to moderate resistance to Cassava Bacterial blight diseases (CBB) developed at the International Institute of Tropical Agriculture (IITA) Nigeria from crosses between elite genotypes from West Africa crossed to genotypes introduced from either East Africa or Latin America (Table 1). Two elite white root, commercially cultivated genotypes were used as checks; TMEB419 (CMD resistant and preferred for starch production) and TMEB693 (CMD resistant with excellent mealiness for food uses). The 26 genotypes were grown in field trials at five locations in Nigeria during the 2014-2015 cropping season. The test locations Mokwa, Ibadan, Ikenne, Ubiaja and Zaria represent the major cassava growing agro ecologies in the country. The agro ecological characteristics of the locations, description of the climatic and soil characteristics are presented in Table 2. The experiment was established using a Randomised complete block design (RCBD) with 4 replications at each location. The test genotypes were planted in a plot of 7 rows with 6 stands on each row using a planting spacing of 1m x 1m. All measurements were based on the inner five rows discarding the first and the last stands on each ridge to give a total of 20 plants per net plot. No fertilizer was applied. Weeds were controlled as necessary. Harvesting at all locations was at 12 months after planting (MAP).

**Table 1.** Genotypes used from the study and their pedigree.

| Entry | Genotype | Pedigree |
| --- | --- | --- |
| G1 | TMS-IBA090454 | 94/0561 (4×) × 06/0813 |
| G2 | TMS-IBA090482 | 91934 × CM6119-5 |
| G3 | TMS-IBA090488 | 97/2205 × FECULA |
| G4 | TMS-IBA090498 | 97/2205 × MAUNJILI |
| G5 | TMS-IBA090504 | 97/2205 × MKONDEZI |
| G6 | TMS-IBA090506 | 97/2205 × MKONDEZI |
| G7 | TMS-IBA090508 | 97/2205 × MKONDEZI |
| G8 | TMS-IBA090509 | 96/1632 × CM 5306-8 |
| G9 | TMS-IBA090510 | 97/2205 × MKONDEZI |
| G10 | TMS-IBA090516 | 97/3200 × KIBAHA |
| G11 | TMS-IBA090520 | 97/4763 × KALESO |
| G12 | TMS-IBA090521 | 97/4763 × MAUNJILI |
| G13 | TMS-IBA090522 | 97/4763 × MAUNJILI |
| G14 | TMS-IBA090523 | 97/4763 × MAUNJILI |
| G15 | TMS-IBA090536 | 96/1569 S1 |
| G16 | TMS-IBA090537 | 96/1569 S1 |
|  |  | 91/02324 × M.COL |
| G17 | TMS-IBA090546 | 1468 |
| G18 | TMS-IBA090564 | 96/1089A × CM6921-3 |
| G19 | TMS-IBA090574 | 97/2205 × MKONDEZI |
| G20 | TMS-IBA090576 | 96/1632 × CM 5306-8 |
|  |  | 96/1632 × M. BRA |
| G21 | TMS-IBA090581 | 1045 |
|  |  | 97/2205 × M. COL |
| G22 | TMS-IBA090590 | 1468 |
| G23 | TMS-IBA090597 | 97/4763 × CM7514-8 |
|  |  | 98/0581 × M. COL |
| G24 | TMS-IBA090609 | 1468 |
| G25 | TMEB419 | Check |
| G26 | TMEB693 | Check |

**Table 2.** Test environments and there climatic features.

| Code | Location | Agro-ecology | Coordinates |
| --- | --- | --- | --- |
| E1 | Ibadan | Forest savannah transition | lat. 7°38'N and lon. 3°89'E |
| E2 | Ikenne | Humid forest zone | lat. 6°86'N and lon. 3°71'E |
| E3 | Ubiaja | sub-humid forest zone | lat. 6°65'N and lon. 6°38'E |
| E4 | Zaria | Northern Guinea savannah | lat. 11°86'N and lon. 7°42'E |
| E5 | Mokwa | Southern Guinea savannah | lat. 9°28'N and lon. 5°05'E |

### Data collection and analysis

At harvest, measurements were made on root number, root weight (fresh root yield) and shoot weight (data not shown). Roots were sampled from each plot across the replications and processed for estimation of dry matter content (DMC) following the oven dry method described by Norbert et al. (2012). Fresh root yield (FYLD) t/ha and the (DMC) per plot was further used to derive the dry yield (DYLD) in tonnes per hectare. The plot means for DYLD were analysed using the analysis of variance (ANOVA) for within and across locations. The data was further subjected to the GEI analysis using the GGE biplot (Yan, 2001; Yan & Kang, 2003; Yan et al., 2007) and Yield selection index (YS_I_) by Kang and Magari (1995).

## Results

Variations due to Genotype, Environment and Genotype x Environment were significant for DYLD (Data not shown).

Overall mean DYLD (t/ha) of the 26 genotypes across locations was 6.31 t ha^−1^, and ranged from 3.63 t ha^−1^ (TMS-IBA090516) to 8.15 t ha^−1^ for TMS-IBA090506 (Table 3). Ibadan had a mean of 9.79 t ha^−1^, and ranged from 2.58 t ha^−1^ (TME419B) to15.45 t ha^−1^ (TMS-IBA090536). 67% of the improved genotypes had yield above the location mean, while the two checks yielded below the improved clones (Table 3). Ikenne had a mean of 7.39 t ha^−1^, with a range of 3.76 t ha^−1^ (TMS-IBA090516) to 10.98 t ha^−1^ (TMS-IBA090574). 62% of the improved genotypes performed above average while 71% outperformed at least one of the checks (Table 3). The mean performance at Mokwa was 5.05t ha^−1^, and ranged from 3.47 t ha^−1^ (TMS-IBA090488) to 6.57 t ha^−1^ (TMS-IBA090506 & TMS-IBA090576). The mean performance at Ubiaja ranged from 2.48 t ha^−1^ (TMEB 693) to 8.49 t ha^−1^ (TMS-IBA090506), with a mean of 5.57 t ha^−1^. 58% of the improved materials had above average performance while 38% out yielded at least one of the commercial checks. Performance in Zaria ranged from 1.52 t ha^−1^ (TMS-IBA090546) to 6.72 t ha^−1^ (TMS-IBA090590) with a mean of 3.73 t ha^−1^. 38% of the improved clones performed above average while 25% outperformed at least one of the checks.

**Table 3.** Mean dry root yield (t/ha) of 26 cassava clones evaluated at Ibadan, Ikenne, Mokwa, Ubiaja and Zaria in 2014-2015.

| clone | Ibadan | Ikenne | Mokwa | Ubiaja | Zaria | Mean±SE |
| --- | --- | --- | --- | --- | --- | --- |
| TMS-IBA090506 | 9.32 | 10.14 | 6.57 | 8.49 | 6.24 | 8.15±0.8 |
| TMS-IBA090581 | 14.4 | 9.97 | 4.17 | 5.4 | 3.72 | 7.53±2 |
| TMS-IBA090590 | 10.86 | 8.41 | 4.93 | 6.69 | 6.72 | 7.52±1 |
| TMS-IBA090509 | 11.93 | 8.17 | 6.15 | 5.22 | 5.12 | 7.32±1.3 |
| TMS-IBA090521 | 11.42 | 9.39 | 3.63 | 6.4 | 5.72 | 7.31±1.4 |
| TMS-IBA090576 | 10.55 | 8.57 | 6.57 | 6.41 | 3.93 | 7.21±1.1 |
| TMS-IBA090574 | 10.75 | 10.98 | 4.16 | 7.65 | 2.33 | 7.17±1.7 |
| TMS-IBA090523 | 11.76 | 5.93 | 6.4 | 6.47 | 4.75 | 7.06±1.2 |
| TMS-IBA090536 | 15.45 | 8.1 | 4.31 | 3.65 | 3.16 | 6.93±2.3 |
| TMS-IBA090597 | 12.27 | 7.49 | 6.02 | 5.01 | 3.7 | 6.9±1.5 |
| TMS-IBA090609 | 11.36 | 7.2 | 4.23 | 6.14 | 5.52 | 6.89±1.2 |
| TMS-IBA090510 | 12.6 | 5.81 | 6.38 | 4.45 | 4.22 | 6.69±1.5 |
| TMS-IBA090454 | 9.49 | 7.96 | 6.5 | 5.28 | 3.47 | 6.54±1 |
| TMS-IBA090504 | 10.29 | 8.75 | 5.55 | 5.1 | 3.01 | 6.54±1.3 |
| TMS-IBA090498 | 8.79 | 5.04 | 5.97 | 6.5 | 5.63 | 6.39±0.6 |
| TMS-IBA090564 | 8.56 | 7.84 | 5.49 | 7.21 | 2.53 | 6.33±1.1 |
| TMS-IBA090488 | 12.87 | 9.16 | 3.47 | 3.54 | 1.87 | 6.18±2.1 |
| TMS-IBA090482 | 9.41 | 7.51 | 5.52 | 4.95 | 3.07 | 6.09±1.1 |
| TMS-IBA090522 | 10.72 | 6.24 | 4.6 | 4.95 | 3.3 | 5.96±1.3 |
| TMS-IBA090537 | 9.52 | 7.13 | 4.33 | 5.06 | 3.01 | 5.81±1.1 |
| TMS-IBA090546 | 5.41 | 8.17 | 3.72 | 7.53 | 1.52 | 5.27±1.2 |
| TMS-IBA090508 | 7.44 | 4.78 | 5.71 | 5.43 | 2.17 | 5.11±0.9 |
| TMS-IBA090520 | 6.24 | 4.68 | 4.39 | 6 | 3.08 | 4.88±0.6 |
| TMEB419 | 2.58 | 4.58 | 3.93 | 6.18 | 5 | 4.45±0.6 |
| TMEB693 | 5.05 | 6.4 | 4.52 | 2.48 | 1.66 | 4.02±0.9 |
| TMS-IBA090516 | 5.37 | 3.76 | 4.01 | 2.56 | 2.43 | 3.63±0.5 |
| Mean | 9.79 | 7.39 | 5.05 | 5.57 | 3.73 | 6.31 |
| CV (%) | 13 | 20 | 26.6 | 17.5 | 28.6 |  |
| LSD(0.05) | 2.53 | 2.95 | 2.67 | 1.95 | 2.12 |  |

The GGE biplots (fig 1, 2 & 3) captured the responses of the twenty six (26) cassava genotypes evaluated for DYLD (t ha^−1^) at five locations. The first and second principal components accounted for 78.1% of the total GGE variation for DYLD. The polygon view of the biplot (also called ‘which won where’) showed that the entire test environments fell into two of the five sectors outlined and identified two mega environments; Ikenne (E2), Ubiaja (E3), Zaria (E4) and Mokwa (E5) on one hand, and Ibadan (E1) formed the other (fig. 1). Ibadan (E1) mega environment had TMS-IBA090536 (15) as the winning genotype. Other genotypes that fell into this mega environment were TMS-IBA090581 (21), TMS-IBA090488 (3), TMS-IBA090510 (9), TMS-IBA090522 (13), TMS-IBA090597 (23), TMS-IBA090504 (5) and TMS-IBA090523 (14). TMS-IBA090506 (6) was the superior genotype in the other mega environment that also contained TMS-IBA090590 (22), TMS-IBA090574 (19), TMS-IBA090521 (12), TMS-IBA090576 (20), TMS-IBA090609 (24), TMS-IBA090564 (18) and TMS-IBA090454 (1) (fig 1). 9 genotypes (TMS-IBA090516 (10), TMEB693 (26), TMS-IBA090508 (7), TMS-IBA090537 (16), TMS-IBA090482 (2), TMS-IBA090520 (11), TMEB419 (25), TMS-IBA090546 (17) and TMS-IBA090498 (4) fell into sectors that contained no test location (fig 1). Fig 2 shows mean performance *versus* stability ranking of the 26 genotypes. The two lines that passed through the origin are the ordinate (blue) and the abscissa (red) of the Average Environment Coordinate (AEC). The AEC abscissa points in the direction of higher mean performance of the genotypes and, consequently ranks the genotypes with respect to mean performance (Yan et al, 2007). The intercept of the AEC abscissa on the ordinate therefore divided the genotypes into high (genotypes on the right) and low yielding (genotypes on the left). Thus, across the five locations genotypes performance for DYLD were ranked as follows for the top seven genotypes: TMS-IBA090506 (6),TMS-IBA090581 (21),TMS-IBA090574 (19), TMS-IBA090521 (12), TMS-IBA090590 (22), TMS-IBA090509 (8), TMS-IBA090576 (20)= TMS-IBA090536 (15). The low yielding clones were rank in decreasing order as follows; TMS-IBA090498 (4), TMS-IBA090482 (2), TMS-IBA090522 (13), TMS-IBA090537 (16), TMS-IBA090546 (17), TMS-IBA090508 (7), TMS-IBA090520 (11), TMEB419 (25), TMEB693 (26) and TMS-IBA090516 (10). The projections on the ordinate, depending on the length, measure the stability of the genotypes. Thus, the shorter the vector the more stable is the associated genotype and vice versa (Yan and Kang, 2003). TMS-IBA090454 (1) had the shortest vector, followed by TMS-IBA090574 (19), TMS-IBA090576 (20), TMS-IBA090482 (2), TMS-IBA090590 (22) and TMS-IBA090521 (12); four of these genotypes had a near zero projection onto the AEC ordinate. TMEB419 (25) had the longest projection on the AEC ordinate. Other genotypes associated with longer vectors were TMS-IBA090536 (15), TMS-IBA090546 (17), TMS-IBA090488 (3), TMS-IBA090581 (21) and TMS-IBA090520 (11). Three of the six stable genotypes (TMS-IBA090574 (19), TMS-IBA090521 (12) and TMS-IBA090590 (22) were among the top seven high yielding genotypes. The small circle close to the arrow of the AEC abscissa delineates the ideal genotype and TMS-IBA090574 (19) was the closest to the circle. Figure 3 shows the discriminatory ability and representativeness of the test locations. The length of the environment vectors (which approximates the standard deviation within each environment) from the biplot origin and the angle formed with the abscissa of the AEC reveals the discriminatory ability and the representativeness of the test locations respectively, (Yan and Kang, 2003). The longer the vector, the higher the discriminatory ability of the associated environment and the shorter the angle formed, the more representative the associated environment (Yan et al., 2007). The biplot identified Ikenne (E2) as the most representative since its vector formed the shortest angle with the AEC abscissa. It was followed by Mokwa (E5) and Zaria (E4) sequentially, while Ubiaja (E3) and Ibadan (E1) with the largest angles were least representative of their mega environment. Ibadan (E1) had the highest discriminatory power due to its possession of the longest vector, followed by Ubiaja (E3), Ikenne (E2) and Zaria (E4). Mokwa (E5) with the shortest vector was the least environment in terms of discriminating power. The small circle close to the arrow of the AEC abscissa delineates the ideal environment (best for cultivar evaluation) and the location closest to it is adjudged the best (Yan and Kang, 2003). Ikenne (E2) was closest to the ideal environment and therefore picked as the best (fig 3).

**Figure 1.**
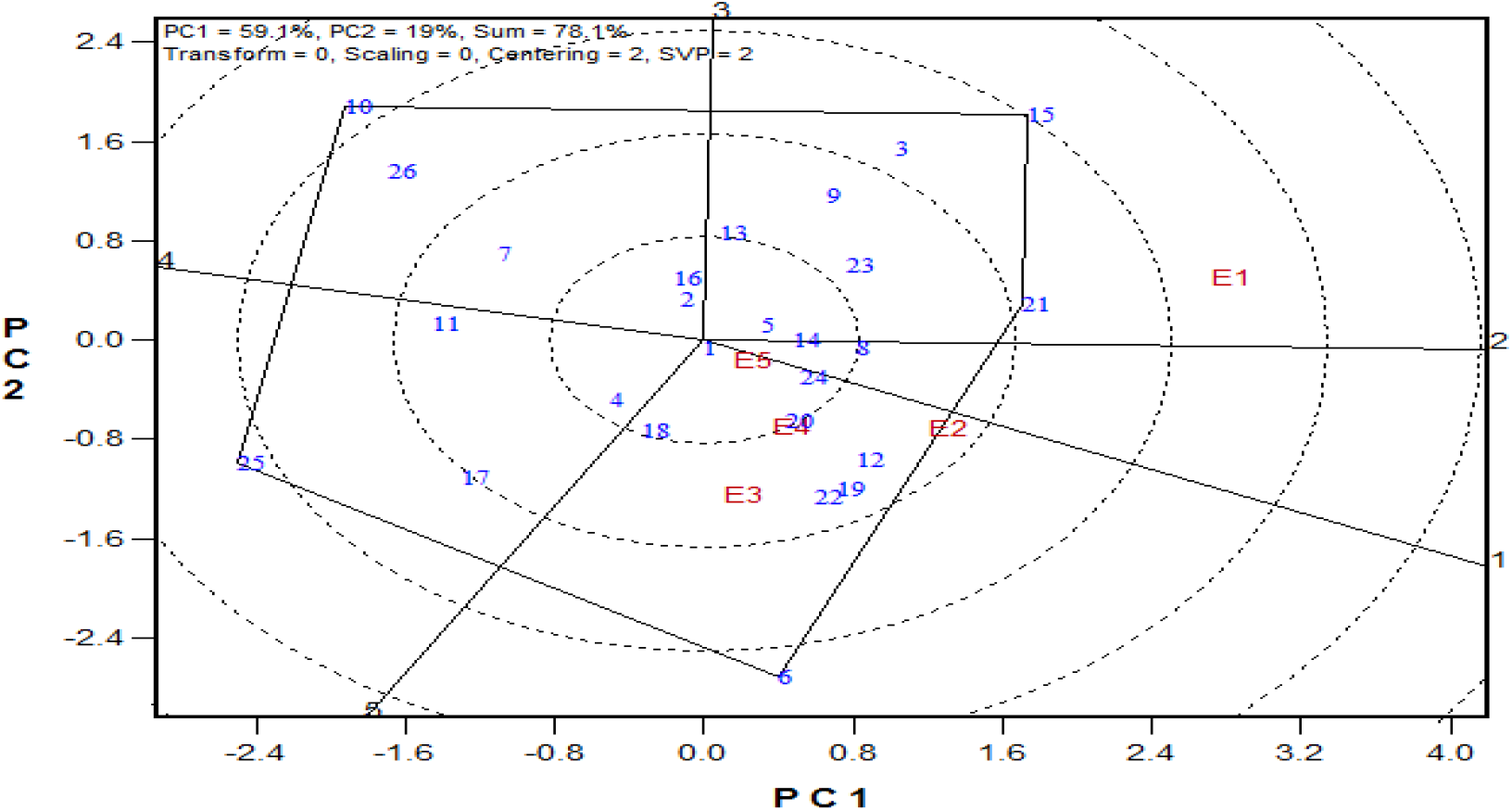
The “which-won-where” view of the GGE biplot for Dry root yield

**Figure 2.**
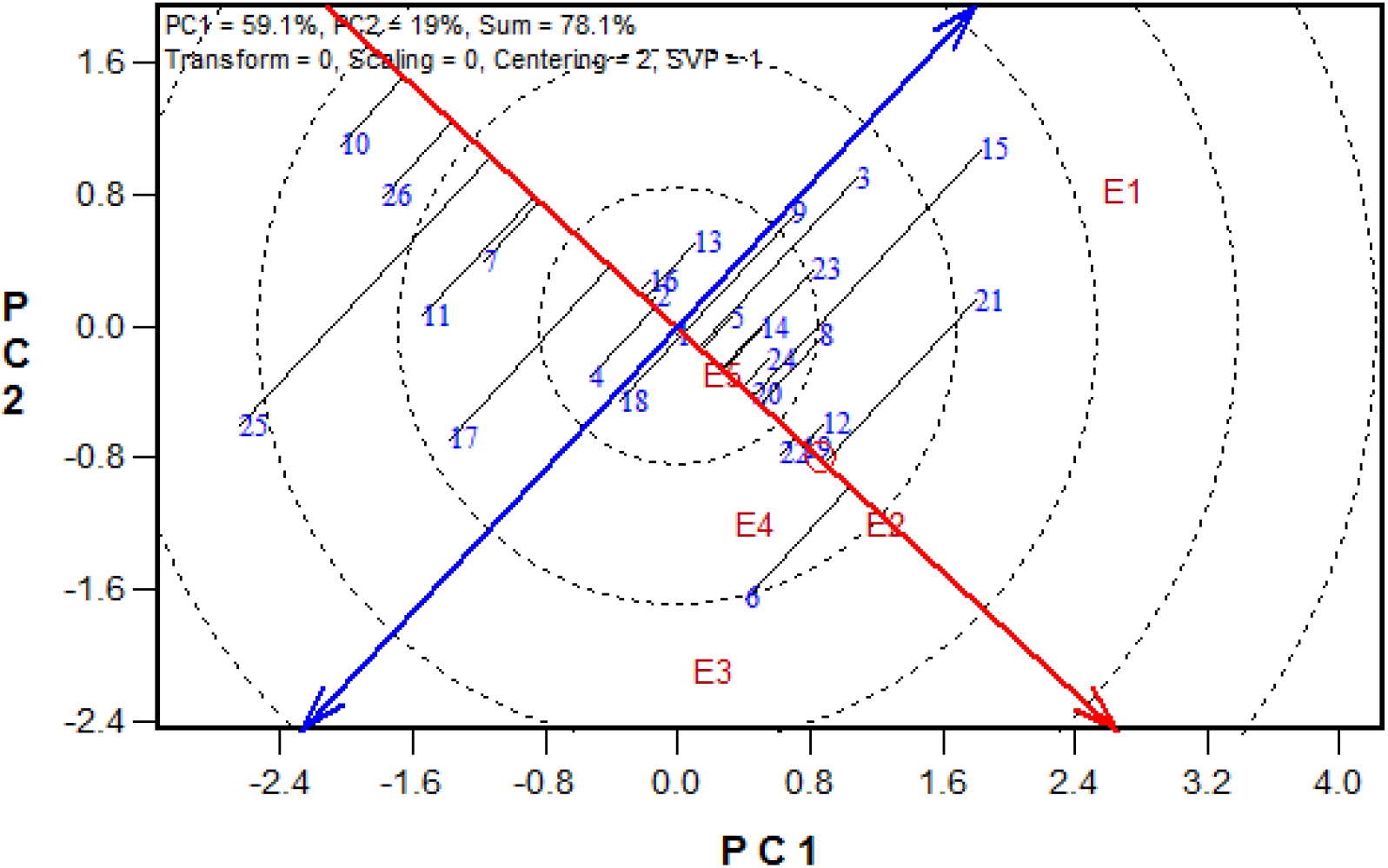
The “mean vs. stability” view of the GGE biplot for Dry root yield

**Figure 3.**
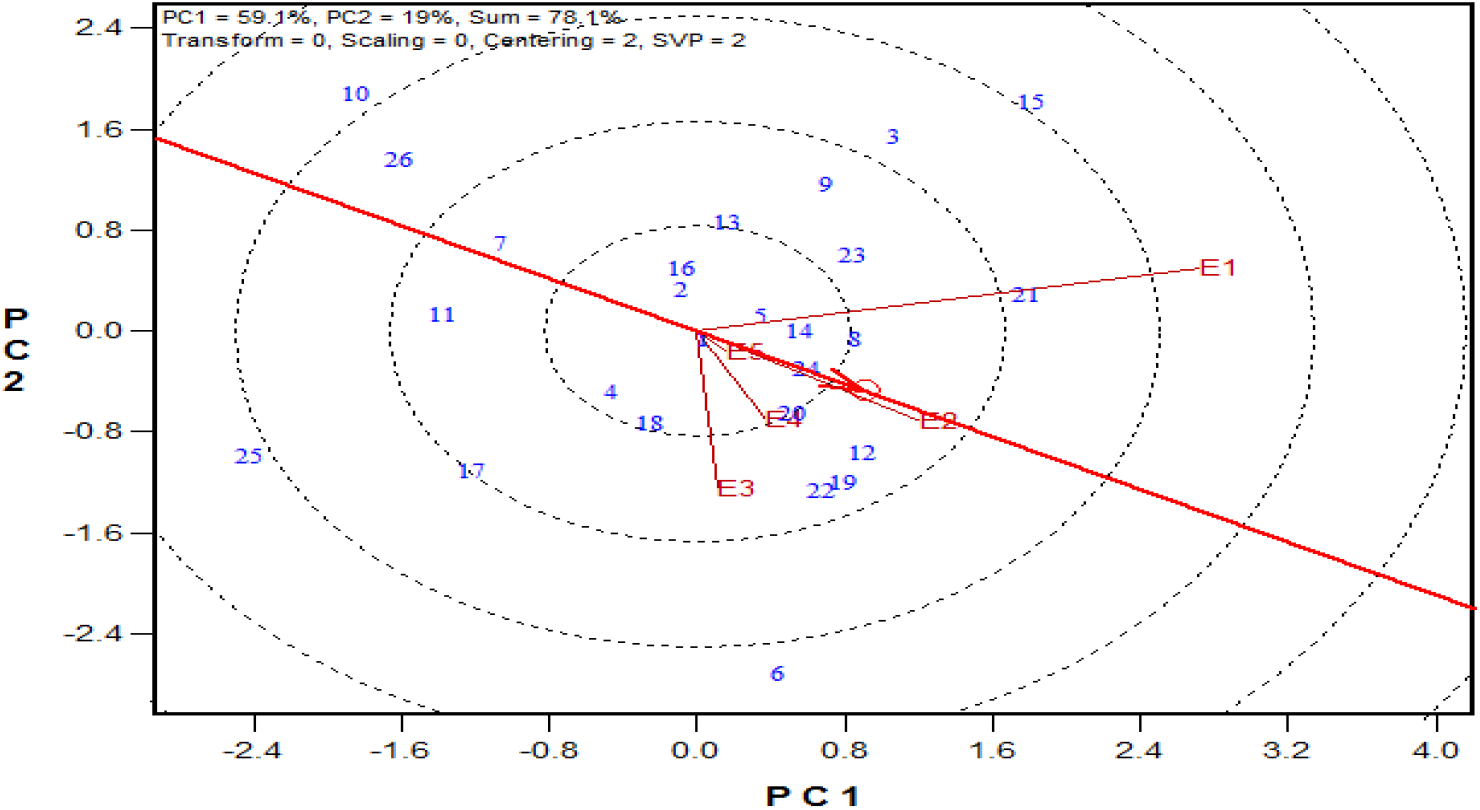
The “discriminating power vs. representativeness” view of the GGE biplot for dry root yield

Table 4 shows the yield, rank and adjusted rank, Shukla stability variance and rating, and YSi of 26 cassava genotypes evaluated for DYLD. TMS-IBA090506 with the highest yield (8.53 t/ha) got the highest rank (26), followed by TMS-IBA090581 (7.53 t/ha) with 25. The lowest yielding genotype across environments, TMS-IBA090516 (3.26 t/ha) had the lowest rank of 1. TMS-IBA090581, TMS-IBA090590, TMS-IBA090521, TMS-IBA090509 and TMS-IBA090506 that were 1 LSD more than mean (6.30 t/ha) got a compensation of 2 which increased their adjusted ranks from 25-27, 24-26, 22-24, 23-25, 26-28, respectively. TMS-IBA090454, TMS-IBA090498, TMS-IBA090504, TMS-IBA090510, TMS-IBA090523, TMS-IBA090536, TMS-IBA090564, TMS-IBA090574, TMS-IBA090576, TMS-IBA090609, TMS-IBA090597 that were just higher than the mean got a compensation of 1 and a shift in the adjusted rank from 13-14, 12-13, 13-14, 15-16, 19-20, 18-19, 11-12, 20-21,21-22, 16-17, 17-18, respectively. TMS-IBA090454, TMS-IBA090482, TMS-IBA090504, TMS-IBA090509, TMS-IBA090522, TMS-IBA090537, TMS-IBA090564, TMS-IBA090576, TMS-IBA090590, TMS-IBA090597 and TMS-IBA090609 were identified as stable genotypes following their non-significant stability variances (Shukla, 1972) and therefore got a rating of 0. TMS-IBA090523 with a 10% significant stability variance was adjudged unstable and got a rating of −2, while TMS-IBA090506, TMS-IBA090508, TMS-IBA090516 and TMS-IBA090521 significant at 5% got a rating of −4. A rating of −8 was allotted to TMS-IBA090488, TMS-IBA090498, TMS-IBA090510, TMS-IBA090520, TMS-IBA090536 and TMS-IBA090546 with stability variances significant at 1%. The YSi technique selects genotypes as high yielding and stable if they are associated with YSi values higher than the YSi mean. Genotypes TMS-IBA090454, TMS-IBA090504, TMS-IBA090506, TMS-IBA090509, TMS-IBA090521, TMS-IBA090523, TMS-IBA090536, TMS-IBA090564, TMS-IBA090574, TMS-IBA090576, TMS-IBA090581, TMS-IBA090590, TMS-IBA090597 and TMS-IBA090609 with YSi values of 14, 14, 24, 25, 20, 18, 11, 12, 13, 22, 19, 26, 18, and 17, respectively, were identified as stable and high yielding having met the criteria.

**Table 4.** Mean yield, rank and adjusted rank, Shuka stability variance and rating, and YSi of 26 cassava genotypes evaluated for dry root yield.

| Clones | Mean<br>Yield<br>(Y) | YIELD<br>RANK<br>(X) | Adj-X=Y' | Y'+<br>x=Y | $\delta^2_i$ | Stability<br>rating<br>(S) | Ysi=<br>Y+S | Selected<br>clones |
| --- | --- | --- | --- | --- | --- | --- | --- | --- |
| TMS-IBA090516 | 3.63 | 1 | -3 | -2 | 8.8 | -4 | -6 |  |
| TMEB693 | 4.02 | 2 | -3 | -1 | 11.9 | -8 | -9 |  |
| TMEB419 | 4.45 | 3 | -2 | 1 | 49.3 | -8 | -7 |  |
| TMS-IBA090520 | 4.88 | 4 | -2 | 2 | 11.2 | -8 | -6 |  |
| TMS-IBA090508 | 5.11 | 5 | -2 | 3 | 8.2 | -4 | -1 |  |
| TMS-IBA090546 | 5.27 | 6 | -2 | 4 | 22.4 | -8 | -4 |  |
| TMS-IBA090537 | 5.81 | 7 | -1 | 6 | -0.2 | 0 | 6 |  |
| TMS-IBA090522 | 5.96 | 8 | -1 | 7 | 2.1 | 0 | 7 |  |
| TMS-IBA090482 | 6.09 | 9 | -1 | 8 | 0.5 | 0 | 8 |  |
| TMS-IBA090488 | 6.18 | 10 | -1 | 9 | 24.0 | -8 | 1 |  |
| TMS-IBA090564 | 6.33 | 11 | 1 | 12 | 6.0 | 0 | 12 | + |
| TMS-IBA090498 | 6.39 | 12 | 1 | 13 | 12.3 | -8 | 5 |  |
| TMS-IBA090454 | 6.54 | 13 | 1 | 14 | 2.1 | 0 | 14 | + |
| TMS-IBA090504 | 6.54 | 13 | 1 | 14 | 2.5 | 0 | 14 | + |
| TMS-IBA090510 | 6.69 | 15 | 1 | 16 | 13.5 | -8 | 8 |  |
| TMS-IBA090609 | 6.89 | 16 | 1 | 17 | 4.9 | 0 | 17 | + |
| TMS-IBA090597 | 6.9 | 17 | 1 | 18 | 5.6 | 0 | 18 | + |
| TMS-IBA090536 | 6.93 | 18 | 1 | 19 | 37.5 | -8 | 11 | + |
| TMS-IBA090523 | 7.06 | 19 | 1 | 20 | 6.9 | -2 | 18 | + |
| TMS-IBA090574 | 7.17 | 20 | 1 | 21 | 18.0 | -8 | 13 | + |
| TMS-IBA090576 | 7.21 | 21 | 1 | 22 | 0.5 | 0 | 22 | + |
| TMS-IBA090521 | 7.31 | 22 | 2 | 24 | 8.4 | -4 | 20 | + |
| TMS-IBA090509 | 7.32 | 23 | 2 | 25 | 3.1 | 0 | 25 | + |
| TMS-IBA090590 | 7.52 | 24 | 2 | 26 | 4.9 | 0 | 26 | + |
| TMS-IBA090581 | 7.53 | 25 | 2 | 27 | 22.4 | -8 | 19 | + |
| TMS-IBA090506 | 8.15 | 26 | 2 | 28 | 8.0 | -4 | 24 | + |
| Mean | 6.31 |  |  |  |  |  | 9.81 | 7.1 |
| LSD(0.05) | 0.95 |  |  |  |  |  |  |  |

## Discussion

Highly significant variation observed for dry root yield among the evaluated genotypes indicates the existence of variability and potentials for improvement. The mean dry matter content observed in our study was comparable to results obtained by several authors (Aina et al., 2009, Norbert et al., 2012 and Akinwale et al., 2011) and therefore gives credence to the need to further increase the genetic gain in dry matter content which is crucial in the improvement of dry root yield. In this study, all but one of the improved genotypes outperformed the commercial checks across locations, while within locations 25-100% of the hybrids were better than the checks for DYLD thereby highlighting the potentials of the genotypes to increase production and productivity of cassava in SSA. In this regard, TMS-IBA090506 gave the highest yield and may be selected for further improvement. It was followed by TMS-IBA090581, TMS-IBA090590, TMS-IBA090509, TMS-IBA090521, TMS-IBA090576, TMS-IBA090574 and TMS-IBA090523, that all yielded above 7 t ha^−1^. In contrast TMS-IBA090516 had lowest yield, followed by the two checks and 7 other genotypes that all performed below average, making them undesirable for selection in terms of yield. However, the significant GEI evidenced by the changes in genotypic performance ranking across locations implies a high difference in the response of the genotypes across the test locations (Haldane, 1946; Fernandez, 1991, Yan et al., 2007) which could be due to the variations in climatic and edaphic features of each location. Since crossover interaction stems from imperfect correlation between genotypic performance and environment (Ariyo, 2009), superiority judgement solely based on yield may not be reliable and could lead to devastating consequences (Kang and Magari, 1995; Olayiwola and Ariyo, 2013). This justifies the need for further analysis to probe into GEI and integrate yield performance with stability across locations for a reliable decision on superiority or desirability of the clones. In this study GGE biplot and YSi were employed for GEI analysis.

An understanding of GEI patterns could enhance judicious allocation of resources and reveal which genotype is adapted to where. GGE biplot identified two mega-environments, Ikenne, Mokwa, Ubiaja and Zaria, as the first and Ibadan the other. TMS-IBA090506 and TMS-IBA090536 were the outstanding clones in the two mega-environments, respectively, which implies that the two genotypes were most responsive and well adapted to the environments. They therefore have potentials for further development in such areas. Since mega-environment is a set of locations that consistently share similar genotype as the best (Yan and Rajcan, 2002) the best among such may represent the other without loss of information thereby reducing the cost of cultivar evaluation. This is particularly reliable when identified mega-environments are repeatable over years (Yan & Tinker, 2006).

A cultivar will be desirable if it consistently gave high yields when evaluated across varied environmental conditions (Yan and kang, 2003; Olayiwola & Ariyo, 2013). The GGE biplot selects genotypes based on relative high yield and stability of performance. Though TMS-IBA090506 and TMS-IBA090581 were the highest yielding genotypes, GGE biplot marked them unstable and therefore rendered them undesirable for selection, Aina et al. 2009; Akinwale et al. 2011 and Peprah et al. 2013 also reported high yielding but unstable genotypes. Only TMS-IBA090574, TMS-IBA090521, TMS-IBA090590, TMS-IBA090576 combined high yield with stability according to the biplot, an indication that they were well buffered and could withstand substantial fluctuations across varied growing conditions. The small circle close to the arrow of the AEC abscissa delineates the ideal genotype and since ideal genotype rarely exist in nature, the closest to the circle is a adjudged the ideal (Yan and Kang, 2003). TMS-IBA090574 being the closest to the small circle was identified as the most superior and therefore the most broadly adapted among the clones evaluated in this study.

Yan (2001) stated the ideal environment was the most discriminatory and representative, and designated at the small circle on the abscissa arrow of the test environment evaluation biplot. The most discriminatory environment is that which gives the best information on genotypic differences (performance) while the most representative is that which is closest to the average environment (Yan, 2001; Yan et al., 2007). Again, the ideal environment does not exist in nature and as Ikenne was the closest to stated point, GGE biplot identified it as the ideal test location in this study. Although Mokwa had good representativeness, it had a poor discriminatory ability which was further evident by the non-significant genotypic effect for the DYLD at the location. Mokwa is therefore not suitable for cultivar evaluation as results obtained may not be reliable. Yan & Tinker (2006) discouraged the use of such environment as they not only give little or no encouragement to good genotypes but also have potentials to upgrade poor ones. Ibadan on the other hand was highly discriminative but with poor representativeness; hence it cannot be used in selecting superior genotypes, but could be useful in identifying unstable genotypes for discard.

The YSi statistic by Kang and Magari (1995) is a summation of the adjusted rank and the stability rating, and thus compensates genotypes for high yield and stability, and penalize genotypes for low yield and being unstable. Genotypes TMS-IBA090454, TMS-IBA090504, TMS-IBA090506, TMS-IBA090509, TMS-IBA090521, TMS-IBA090523, TMS-IBA090536, TMS-IBA090564, TMS-IBA090574, TMS-IBA090576, TMS-IBA090581, TMS-IBA090590, TMS-IBA090597 and TMS-IBA090609 were considered to be high yielding and stable. Six of these genotypes (TMS-IBA090523, TMS-IBA090536, TMS-IBA090574, TMS-IBA090581, TMS-IBA090506, TMS-IBA090521) were actually marked as unstable by the Shukla’s stability variance which indicates a weakness for the YSi. It appears that the YSi gave premium to high yield rank which could mask negative stability rating thereby paving the way for the selection of unstable genotypes. A genotype could get high yield rank even if it was not fantastic at all locations. This is particularly true for TMS-IBA090536 that performed above average at only two of the tested sites. The selection of high yielding but unstable genotypes does not have any practical advantage and may lead to commercial losses (Kang & Magari, 1995; Waldron et al., 2002; Olayiwola & Ariyo, 2013). It becomes imperative to employ YSi with caution when genotype with broad adaptation is the target. It is a usual practice for breeders to compare statistical techniques for an efficient and reliable decision in breeding programs. A comparison of the selection for desirability made by the GEI techniques employed in this study showed some agreements. The two techniques identified TMS-IBA090590 and TMS-IBA090576 as high yielding and stable genotypes. These two clones performed above average in at least four of the five locations used for evaluation. Yan & Kang (2003) and Nassir & Ariyo (2011) found selection agreement between both techniques in independent studies on wheat and rice, respectively. These two genotypes may therefore be considered for further improvement and release for increased cassava yield in Nigeria.

